# BEYOND UNIFORMITY: Pyomelanin’s structural complexity impacts on UV shielding in *Pseudomonas* species with different lifestyles

**DOI:** 10.1101/2024.04.11.589128

**Authors:** Mateo N. Diaz Appella, Adriana Kolender, Oscar J. Oppezzo, Nancy I. López, Paula M. Tribelli

## Abstract

Melanin, a polymeric pigment synthesized by various organisms, confers advantageous traits, including heightened resistance to stress agents. In *Pseudomonas*, disruption of the tyrosine degradation pathway leads to pyomelanin production. Despite a shared synthetic pathway, the chemical structure of pyomelanin remains elusive due to its heterogeneous polymeric nature, suggesting composition variations even among closely related species. Our objective was to analyze pyomelanin structural features across *Pseudomonas* strains: CRISPR/nCas9-engineered *hmgA* mutants of PAO1 and PA14, reference strains of the human opportunist pathogen *P. aeruginosa*; a natural melanogenic mutant (PAM) from a patient; and a Tn5 mutant of the extremophile bacterium *P. extremaustralis* (PexM). Structural analysis revealed strain-specific differences. UV spectra exhibited dual peaks for PAO1 and PA14 mutants, while PAM and PexM displayed a single peak. FTIR indicated changes in the alcohol content ratio, with PAO1 and PA14 *hmgA* mutants having a near 1:1 ratio, PexM a dominant phenol band, and PAM a predominance of the alcohol band. Complex NMR spectra suggested non-linear polymers composed by substituted phenolic rings, carboxylic acids, and alkyl chains, highlighting inter-pigment disparities. UVC (254 nm) susceptibility assessment showed increased survival with pyomelanin addition, correlating with the attenuation of the incoming radiation due to absorption in the culture medium. Moreover, survival to UVC of *P. extremaustralis* was different depending on the melanin source being the most protective pyomelanin obtained from PAO1. These findings reveal distinct pyomelanin subgroups based on structure among strains, elucidating varied physiological effects.

## Introduction

Melanins are polymeric brown pigments widely produced in nature mainly associated with tyrosine metabolism. The general process includes the oxidation of hydroxylated aromatic compounds to quinones that afterwards polymerize to form these heteropolymeric pigments, resulting in different types of melanins (1). Among them, eumelanin or DOPA-melanins are produced by both eukaryotic and bacterial cells through the activity of tyrosinase or laccase enzymes, converting tyrosine to dopaquinone. Several bacterial species mainly produce pyomelanin, named after its discovery in *Pseudomonas aeruginosa* (2). Pyomelanin derives from the accumulation of homogentisate during tyrosine catabolism. Homogentisate formation includes an initial deamination and the complex action of the enzyme 4 -hydroxyphenylpyruvate dioxygenase, encoded by *hppD*. The melanin is formed when homogentisate is accumulated due to an increment of activity of the HppD enzyme or a mutation in the coding gene of the downstream enzyme HmgA. In wild type strains, homogentisate is converted by different enzymes in fumarate and acetoacetate as final products which get into central metabolism. Alterations in the homogentisate pathway leading to spontaneous production of pyomelanin are frequently observed in nature (3). It has been found that melanins confer adaptive advantages by increasing fitness under stressful conditions of bacteria having different lifestyles (4). For example, the production of melanin enhance survival under harsh conditions, such as fluctuations in oxygen levels or the presence of pollutants (5–7). Moreover, blocking the pyomelanin production in *P. aeruginosa* clinical isolates lead to an increase in oxidative stress sensitivity and in *P. aeruginosa* reference strains pyomelanin production provokes a decrease in the clearance in lungs (8). In *Vibrio cholerae* a pyomelanogenic strain showed higher ability to colonize mice intestines (9). Due to their attractive properties and biotechnological interest, melanins have been widely studied (10). Concerning its chemical structure, pyomelanin is a type of nitrogen-free melanin with a proposed linear polymeric structure, keeping intact the −CH_2_COOH group from homogentisate after polymerization, and coupling through the ortho position of the phenol (10). However, there is a lack of accurate description of the complete structure.

The aim of this study was to analyze structural features of pyomelanin from *Pseudomonas* species with diverse lifestyles. We analyzed chemical characteristics as well as the physiological role of pyomelanin by analyzing UVC radiation tolerance of melanogenic strains of the human opportunistic pathogen, *P. aeruginosa* and *P. extremaustralis* an extremophile bacterium from Antarctica. We hypothesized that pyomelanin composition varies across even closely related species conferring distinct biological roles to bacterial producers. Results open interesting avenues for both pathogen control and other biotechnological applications.

## Results

### Pyomelanin production in *Pseudomonas* species with different lifestyles

To study the pyomelanin production and the pigment characteristics in the selected *Pseudomonas* species we constructed different melanogenic mutant strains. Using the CRISPR/nCAS9 system we introduced a premature stop codon in the *hmgA* gene in two *P. aeruginosa* reference strains, PAO1 and PA14. Additionally, a *P. extremaustralis* 14-3b melanogenic strain was obtained from a transposon random insertion library, further named PexM. Sequencing analysis allowed us to determine that the mini-Tn5 in PexM was inserted in the gene (PE143B_0105645) encoding a putative diguanylate cyclase (Fig. S1) that has not been previously reported to be involved in melanin synthesis. Lastly, a spontaneous mutant isolated from a patient with an acute obstructive pulmonary disease and untreated asthma was identified by phenotypic characterization followed by PCR (Fig. S2) as a *P. aeruginosa* strain, different of PA14, and further named PAM. The genomic zone of genes related to melanin biosynthesis was compared in all sequenced strains (PAO1, PA14 and 14-3b) showing a conserved region *hmgA* (Fig. S3). All strains showed a conserved copy of *hppD* and presented a second gene encoding the protein QuiC that contains a domain encoding a 3-dehydroshikimate dehydratase and other domain HppD, but experimental evidence showed that was not related to melanin production (11).

The obtained mutant strains showed a typical brown pigmentation expected for melanin production (Fig. S4). Growth was monitored through viable counts (CFU.mL^-1^) for the wild type and the mutant strains in LB medium under aerobic conditions. No differences in the CFU.mL^-1^ were observed for the different *P. aeruginosa* strains while PexM also showed similar growth in comparison with the wild type strain but a significant decrease of CFU.mL^-1^ value (t-test, P<0.05) was observed after 24h culture (Fig. 1A).

**Figure 1.**
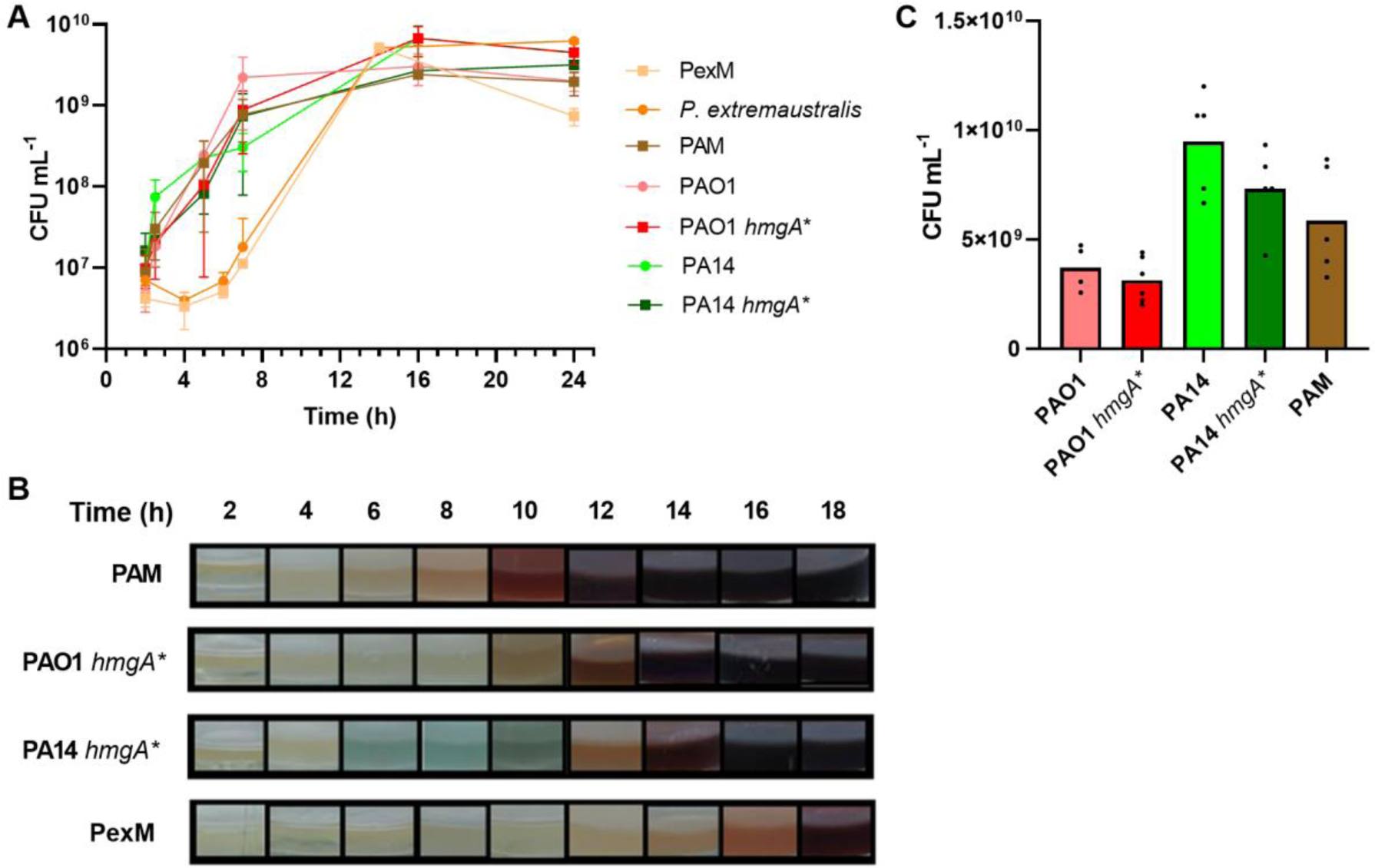
Growth of melanogenic and non-melanogenic strains in different culture media. a: Growth curves of all strains in LB medium. b: Coloration of melanogenic strain in LB cultures through time. c: Growth after 24 h of P. aeruginosa strains in artificial sputum medium (ASM). In (a) values represent mean ± SD of triplicate independent cultures. In (b) bars represent mean and individual measurements are shown.

For *P. aeruginosa* strains, growth was also analyzed in artificial sputum medium (ASM, Sriramulu et al., 2006) that mimic cystic fibrosis lung environment, after 24h of culture. Curiously, growth of PA14 was higher than PAO1, for both wild type and melanogenic strains in this medium (Fig. 1B). PAM strain showed no differences of growth in comparison with reference strains (Fig. 1B).

The analysis of pigment production was conducted in LB medium for all strains, with additional evaluation in ASM medium for *P. aeruginosa* strains. Both PAO *hmgA\** and PA14 *hmgA\** initiated melanin production at approximately 12 h, coinciding with the onset of the stationary phase. Conversely, PexM exhibited a delayed onset of pigment production (16 h), which nonetheless, also corresponded to the stationary phase (Fig. 1B). Notably, PAM presented a different pattern by exhibiting melanin production during the exponential growth phase (Fig. 1B). In terms of concentration, PAM showed the highest melanin concentration reaching 2.10 ± 0.23 mg mL^-1^, while in ASM medium, PA14 *hmgA*\* exhibited the highest melanin production at 1.86 ± 0.21 mg. mL^-1^. Relative melanin production values were comparable across the different strains (Table 1).

**Table 1.** Melanin production in *Pseudomonas* strains after 24 h culture in LB and ASM medium. Mean and standard deviation are showed (n=3). *mg of melanin is relativized to 10^6^ CFU.

| Culture medium | Melanin producing-strain | Net production<br>(mg. mL<sup>-1</sup>) | Relative production*<br>(mg/10<sup>6</sup>CFU) |
| --- | --- | --- | --- |
| LB | PAO1 *hmgA* * | 1.57 ± 0.15 | 3.67*10<sup>-4</sup> ± 1.11*10<sup>-4</sup> |
|  | PA14 *hmgA* * | 1.93 ± 0.15 | 1.67*10<sup>-3</sup> ± 1.87*10<sup>-3</sup> |
|  | PAM | 2.1 ± 0.23 | 1.08*10<sup>-3</sup> ± 4.02*10<sup>-4</sup> |
|  | PexM | 1.27 ± 0.11 | 1.81*10<sup>-3</sup> ± 5.89*10<sup>-4</sup> |
| ASM | PAO1 *hmgA* * | 1.13 ± 0.25 | 3.95 ± 1.46 |
|  | PA14 *hmgA* * | 1.96 ± 0.21 | 5.30 ± 2.37 |
|  | PAM | 1.23 ± 0.15 | 2.44 ± 1.71 |

Melanin production in ASM medium was further examined for the *P. aeruginosa* strains, revealing that PA14 *hmgA*\* consistently displayed the highest relative melanin production, followed by PAO1 *hmgA*\* (Table 1).

To verify that the pigment production was due to the homogentisate accumulation we used bicyclopyrone that inhibits the HppD enzyme. All melanogenic tested strains showed an inhibition of melanin production confirming the involvement of the homogentisate pathway. All strains showed strong inhibition at 0.1 mM concentration after 48 h of incubation (Fig. 2).

**Figure 2.**
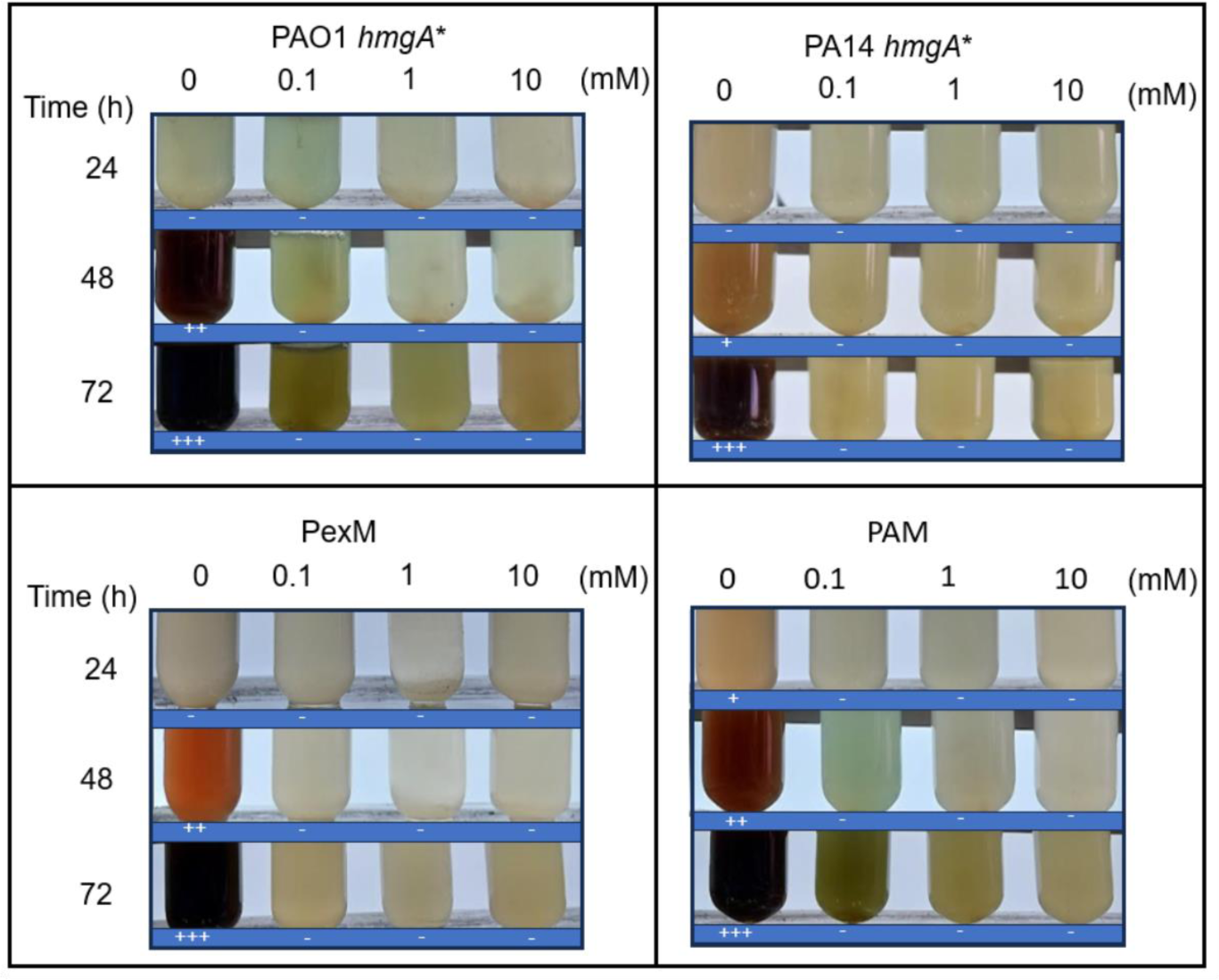
Melanin production of the analyzed *Pseudomonas* strains assayed in LB medium with increasing concentrations of bicyclopyrone. Plus (+) and minus (-) signs indicate the presence or absence of melanin, respectively. An increase of plus (+) signs indicates higher production. The cell growth was similar in all concentrations.

### Structural analysis of melanin from different *Pseudomonas* species

The purified pigments from the different *Pseudomonas* species were subjected to structural analysis including ultraviolet (UV), infrared (FTIR), and ^1^H and ^13^C (NaOD-D_2_O) nuclear magnetic resonance (NMR) spectroscopies.

The UV-Vis spectra of the pigment extracted of the mutant model strains (PAO1 *hmgA*\* and PA14 *hmgA*\*) were very similar, with absorption maxima at 222 and 254 nm of comparable intensities and a minor broad band at 340 nm, while PAM and PexM exhibited the intense maximum only at 222 nm, together with the band at 340 nm suggesting a more extended conjugation system and various oxidation states of the monomeric units constituting the backbone (12) (Fig. 3A). These absorbances agreed with the complex conjugated systems found in melanin structure (13), while the lack of absorption bands in the range 260-280 nm suggested the absence of nucleic acid, lipid, or protein impurities.

**Figure 3.**
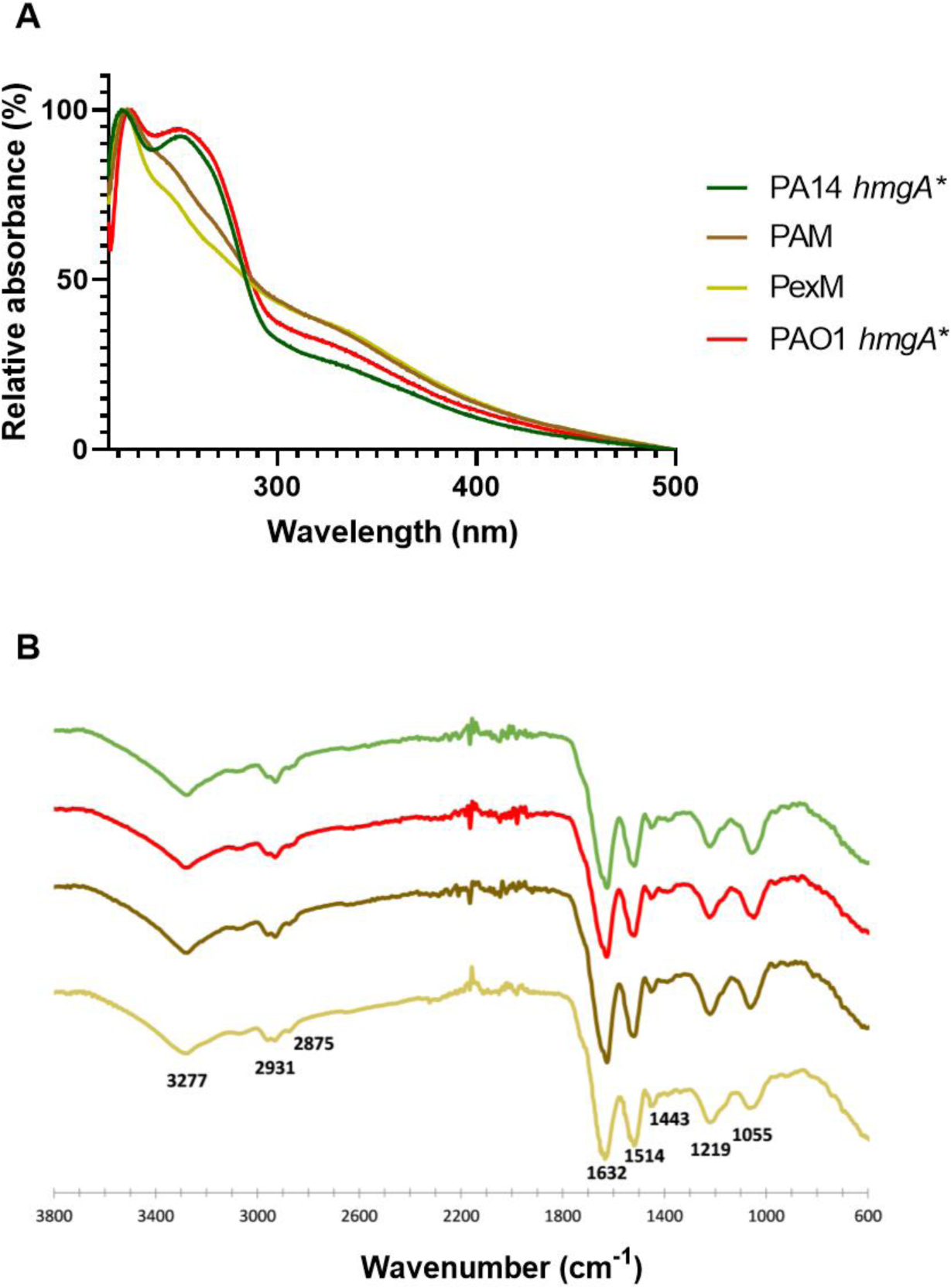
Structural analysis of melanin from different *Pseudomonas* strains: (a) UV-Vis spectra (200-500 nm); (b) FT-IR spectra (ATR, 38000-600 cm^-1^)

FTIR spectra were similar for all strains, with absorptions at 3277 (O−H from carboxylic acid, phenol, alcohol), 2931 (C−H from aromatic moieties), 2875 (C−H associated to *sp*^3^ C), 1632 (C=O from carboxylate), 1514 (aromatic C=C), 1443 (O−H), 1219 (C−O in phenol or phenol ether) and 1055 cm^-1^ (C−O in primary alcohol or ether). Some differences were observed, arising from the relative intensities at 1219 (Ph-O) and 1055 (C-O alcohol, ether) cm^-1^. Thus, PAO1 *hmgA*\*and PA14 *hmgA*\* had 1:1 ratio; while in PexM, the phenol band predominated, and in PAM, the alcohol band. After being subjected to alkaline digestion, a notable decrease in signal intensity at 1055cm^-1^ was observed probably due to etherification or esterification of the associated free hydroxy group (Fig. 3B).

The ^1^H NMR spectra exhibited main broad signals that were consistent with the presence of a polymeric sample. Resonances at 6.4, 6.8 and 7.2 ppm were assigned to aromatic protons included in phenolic rings, while signals in the range 3.5-4.5 ppm indicated alkyl groups (CH, CH_2_ or CH_3_) linked to either N or O, and the range 0.5-3.0 was attributed to alkyl groups (Fig. 4A). On the other hand, the ^13^C NMR spectra showed carbonyl resonances at 170-180 ppm that were assigned to carboxylic acids and esters (Fig. 4B). The region at 140-160 ppm exhibited only weak signals associated to quaternary C−O of phenol residues, while the range 115-145 ppm included more intense resonances of phenolic C−H. Interestingly, melanin from PAO1 *hmgA*\* and PA14 *hmgA*\* showed signals at 60-110 ppm that suggested the presence of carbohydrate moieties (Fig. 4B). At higher field, peaks at 50-60 ppm were associated to CH_2_O, CH_3_O or CH_2_N groups, while the more intense peaks at 10-40 ppm corresponded to alkyl residues (Fig. 4B). All these spectroscopic results agreed with the solubility behavior of the melanin samples, since carboxylic acids and phenols would be deprotonated in alkaline conditions enhancing the solubility of the polymeric samples.

**Figure 4.**
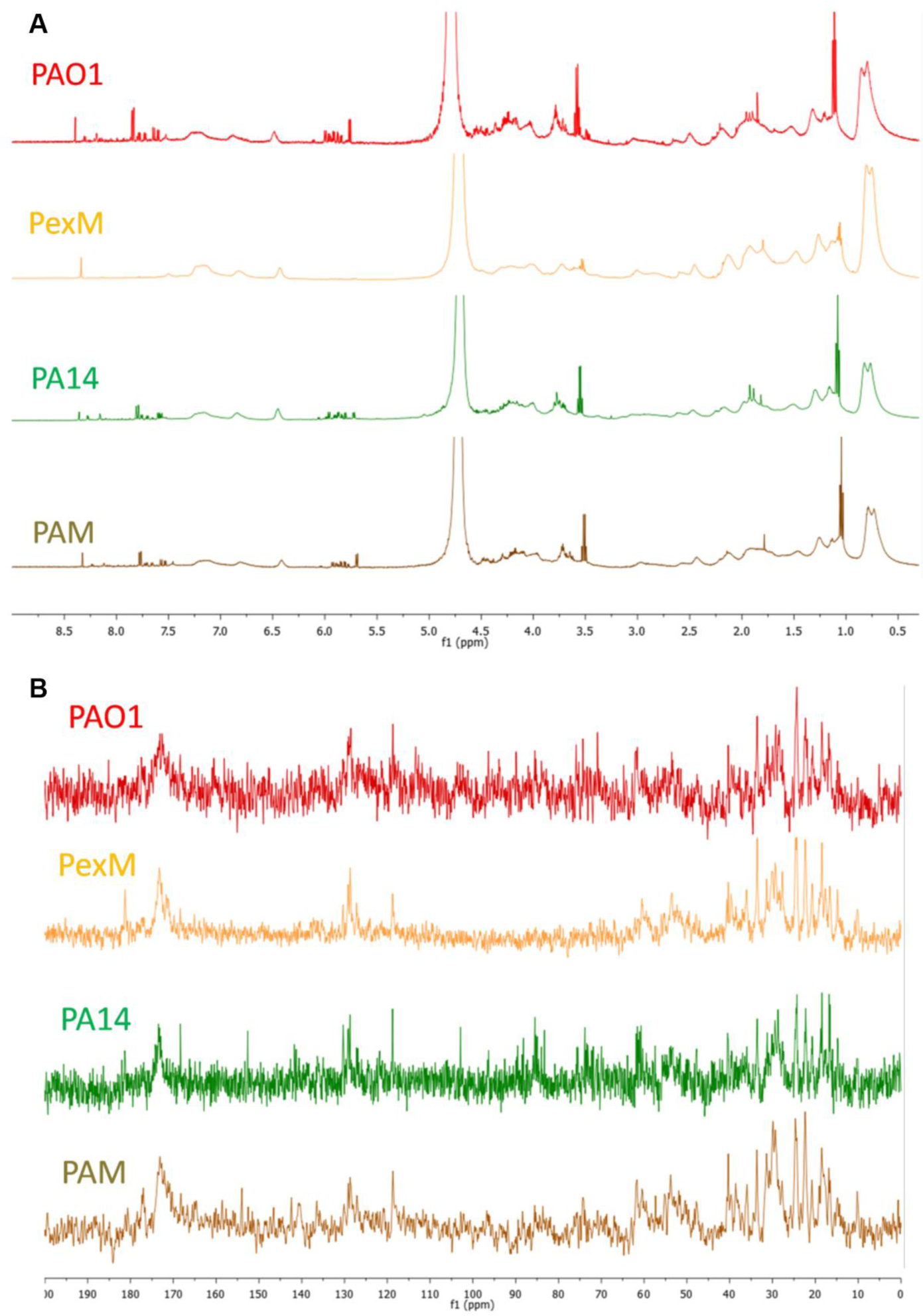
NMR spectra of melanin samples isolated from different strains. A. ^1^H NMR (500 MHz, NaOD-D_2_O). B. ^13^C NMR (125.5 MHz, NaOD-D_2_O).

In order to have a quantitative analysis on the melanin composition, three regions of the ^1^H NMR spectra were integrated (Fig. S5-S9) for each sample: (1) the Ar region (7.6-6.3 ppm), corresponding to aromatic protons and given an arbitrary area of 1.00; (2) the RO region (4.6-3.4 ppm), associated to alkoxy moieties; and (3) the aliphatic region (2.8-0.4 ppm), attributed to alkyl residues (Table 2). The general pattern of the spectra was similar; however different ratios were detected for the three NMR regions that could explain the distinct behavior among the strains. Thus, the lowest RO: Ar ratio (1.24: 1.00) corresponded to PexM, in comparison with a 3.65: 1.00 ratio for PAO1 *hmgA*\*. The PexM ratio indicated a low content of aliphatic alcohol and ether moieties, in agreement with the FTIR spectrum; while the high abundance of Ar units could allow a more extended conjugated system indicated by a higher absorption at 340 nm and a lower one at 254 nm in the UV-vis spectrum. On the other hand, PAM strain showed the highest aliphatic: Ar ratio of the NMR spectra being 11.34: 1.00, which implies a very low content of aromatic units in agreement with the FTIR result. This observation was also consistent with the lowest absorption at 254 nm in the UV-vis spectra.

**Table 2:**
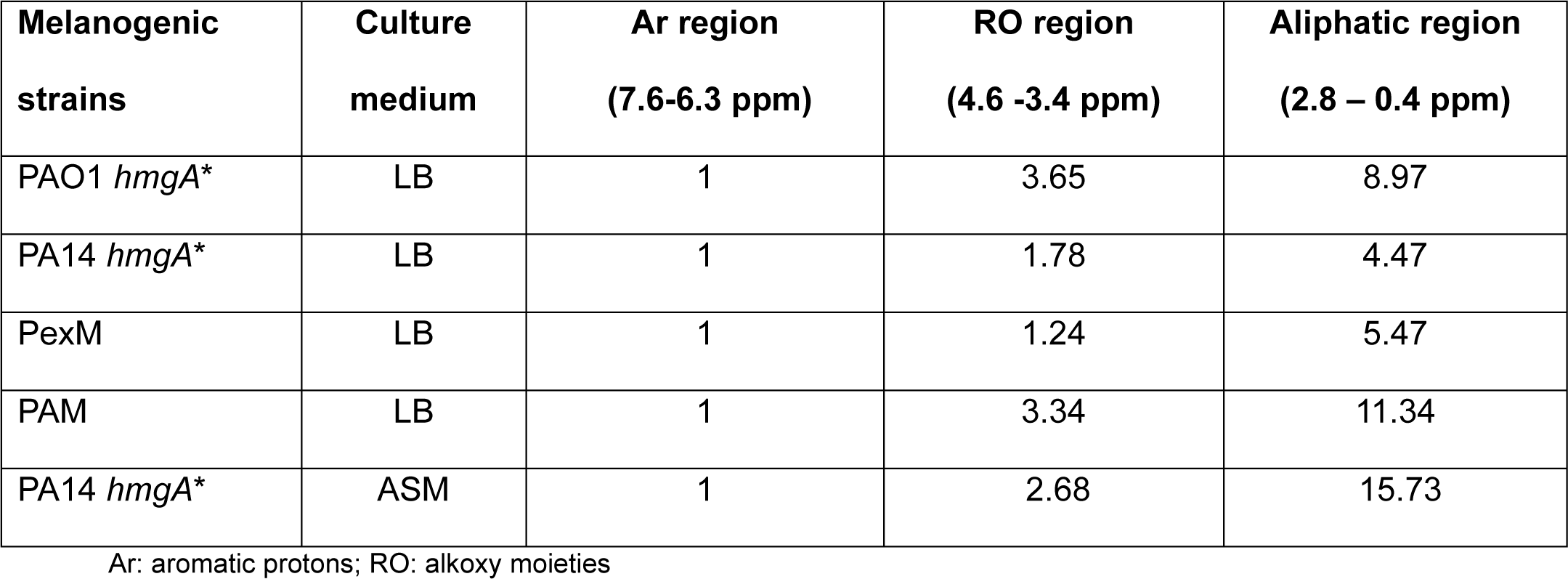
Ratio of the different areas obtained from an ^1^H NMR analysis.

| **Melanogenic strains** | **Culture medium** | **Ar region**<br>**(7.6-6.3 ppm)** | **RO region**<br>**(4.6 -3.4 ppm)** | **Aliphatic region**<br>**(2.8 – 0.4 ppm)** |
| --- | --- | --- | --- | --- |
| PAO1 *hmgA* * | LB | 1 | 3.65 | 8.97 |
| PA14 *hmgA* * | LB | 1 | 1.78 | 4.47 |
| PexM | LB | 1 | 1.24 | 5.47 |
| PAM | LB | 1 | 3.34 | 11.34 |
| PA14 *hmgA* * | ASM | 1 | 2.68 | 15.73 |

Ar: aromatic protons; RO: alkoxy moieties

Comparison between PA14 *hmgA\** grown in LB and ASM revealed a similar general pattern, particularly in the Ar and RO regions, although the RO: Ar ratio was higher for the ASM culture (2.68:1.00) than the LB medium (1.78:1.00). Additionally, the aliphatic region of the ASM spectrum exhibited the most intense broad signals (Table 2).

Together with the broad signals attributed to pyomelanin, some sharp resonances were detected in the NMR spectra and suggested the presence of heterocycles and unsaturated groups. These residues would be covalently linked to the polymer if the purification procedure of the samples was thorough.

In summary, these analyses allowed the differentiation of two groups of melanins based on their structural similarity: the first group corresponded to reference strains (PAO1 *hmgA*\*and PA14 *hmgA*\*), and the second one to the clinical isolate (PAM) and the environmental strain (PexM). The chemical diversity could be associated with the biological function and biotechnological applications of melanins in different microorganisms.

### Bacterial survival upon exposure to ultraviolet radiation (UVC)

Melanins are known to protect against several stress factors including UV radiation. To investigate if the melanin obtained from the different strains exhibited different UVC absorption capabilities, their absorbance at 254 nm was first determined. The purified pigments showed different absorbance at 254 nm when solubilized in PBS, being higher in the reference mutant strains (2.22 and 2.28 for PAO1 *hmgA*\* and PA14 *hmgA*\*, respectively), followed by PAM (2.02) and lower for PexM (1.74). These results were according to the UV-visible spectra analysis that showed higher absorption in the UV region for the melanin from PAO1 *hmgA*\* and PA14 *hmgA*\* (Fig. 3A).

Next, UVC (254 nm) tolerance was assessed by irradiating samples of wild type and melanogenic mutant strains adjusted to the same cell concentration (OD_600nm=_0.3), with or without the addition of purified melanin. Cell samples were exposed to a fluence rate of 0.93 W.m^-2^ for 135 s without melanin and 360 s with melanin. PAO1 and PA14 *hmgA*\*mutants behaved similarly in agreement with the similar pigment structure. The addition of pyomelanin provoked the displacement of their survival curves in a way comparable to that expected for a reduction in the fluence rate of the incoming radiation of 59 and 58%, respectively. However, in PAM and PexM the displacement was smoother, comparable to the effect of a reduction of 55% and 51% in the fluence rate, respectively. The protective effect of pyomelanin was reflected in higher viable cell counts for all strains when compared with the control suspensions without the addition of this pigment (Fig. 5).

**Figure 5.**
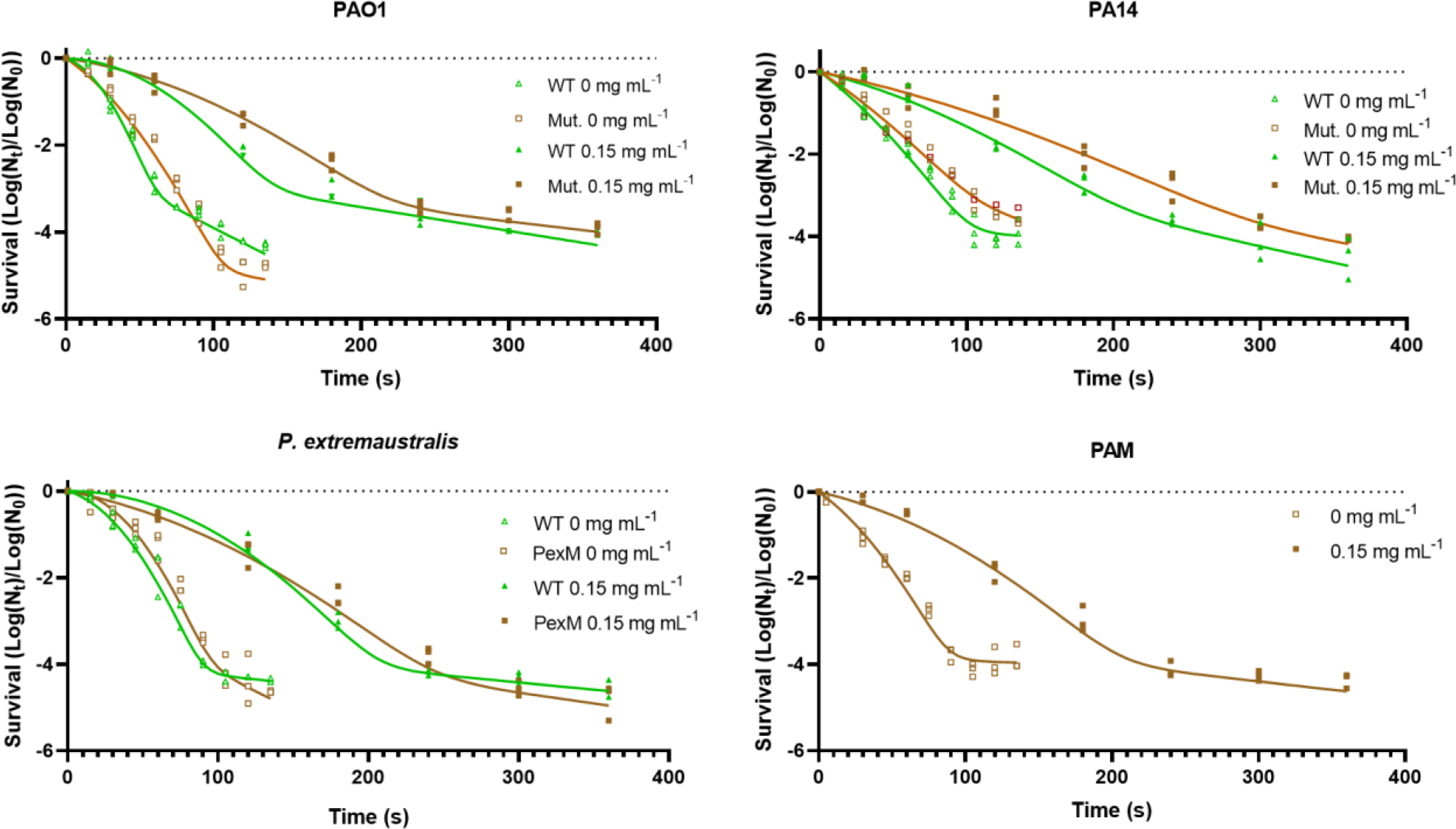
Cell viability after irradiation at 254 nm. The fluency rate dose was constant, equal to 0.93 W m^-2^. Lines represent the functions obtained by fitting the parameters of Eq. 1, showed in Material and Methods section, to the experimental data from assays performed in triplicate (Table S1). WT indicates the wild type strains, and Mut. are its isogenic *hmgA*\* mutants.

We deeply analyzed the different protective effect of melanin by analyzing *P. extremaustralis* survival after 120 s of irradiation without melanin (control) or with the addition of 0.15 mg. mL^-1^ of melanin of the different strains evaluated in this work. *P. extremaustralis* survival showed differences depending on which melanin was added as protector (Fig. 6). In line with the experiments described above, the highest UVC protection was observed for *P. extremaustralis* in presence of PAO1 melanin which also presented the highest absorbance at 254 nm, followed by the PA14 melanin and PAM. The lowest protection was that produced by PexM melanin which showed the lower absorbance at 254 nm.

**Figure 6.**
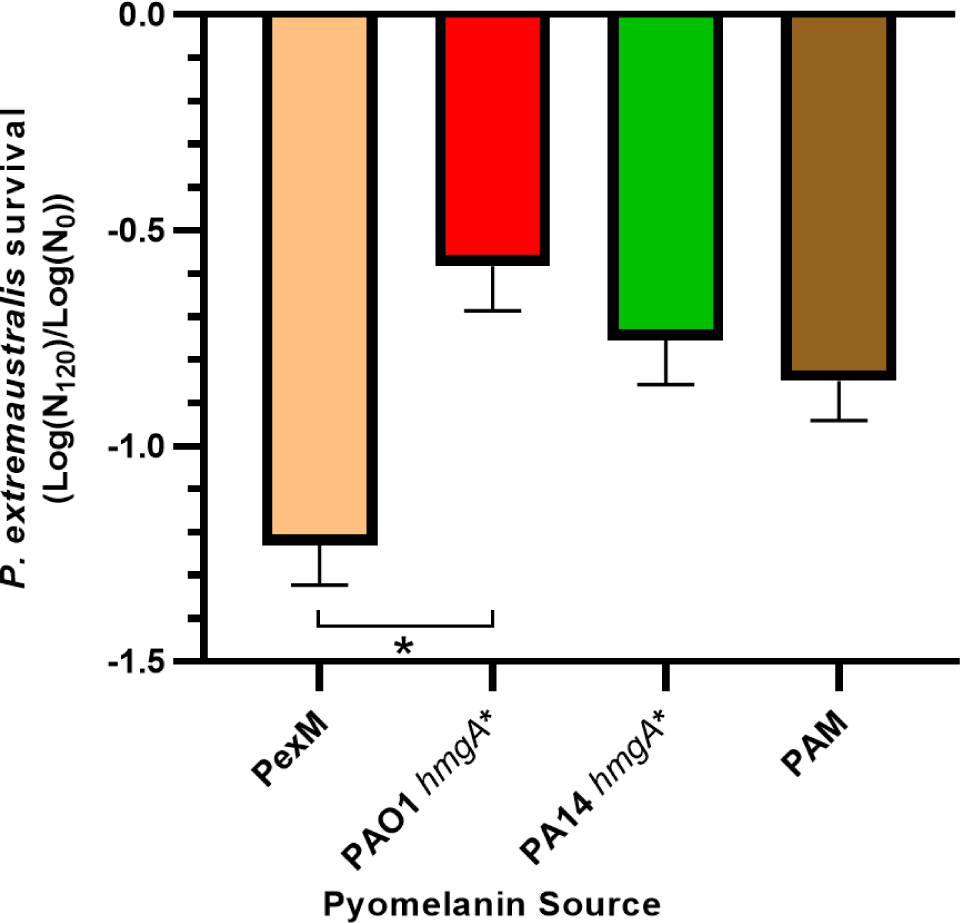
*P. extremaustralis* wild type survival (relative CFU survival) when added 0.15 mg mL^-1^ of melanin extracted from different strains after a 120 s irradiation at 254 nm. The fluency rate dose was constant, equal to 0.93 m W m^-2^

These findings are consistent with the clustering observed in the structures of pyomelanins, with *P. aeruginosa* PAO1 and PA14 demonstrating superior protection and showing that differences in their structure leads to different biological function.

## Discussion

Several living organisms can produce different kinds of melanins, a group of heterogeneous polymeric pigments with relevant biological properties (14). In bacteria, particularly in *Pseudomonas* species spontaneous pyomelanin production, derived of the alteration in the homogentisate pathway, was only reported for *P. aeruginosa* (2,15) and *P. stutzeri* (16). In another *Pseudomonas* species, DOPA melanin is produced by specific enzymes due to the orthohydroxylation of tyrosine to 3,4-dihydroxyphenylalanine (DOPA) and the subsequent transformations that end in the polymerization (3). The structure of melanin is not unique due to its heterogeneous nature (17).

Biotechnological usages have been reported for both types of melanin, those produced by L-DOPA pathway and pyomelanin. For example, melanin is used for the preparation of cosmetics with sun protection factor (SPF) that involve absorbing radiation to prevent photo-induced skin damage as antioxidants, antitumor, and anti-inflammatory compounds amount other uses (18).

In this study, we conducted a comparative analysis of the pyomelanin characteristics produced by two *Pseudomonas* species exhibiting distinct lifestyles. Additionally, we explored the physiological features to ascertain potential variations in the structure of pyomelanins among different strains and their potential impact on functionality.

Melanin has interesting properties, among them the ability to absorb both visible light and UV radiation and in the X-ray range (19). Due to its biotechnological and ecological roles, structural studies have been performed with melanins from different organisms showing heterogeneity (13). However, in our best knowledge there are no studies comparing the structure and biological function of pyomelanins in different bacterial species belonging to the same genus.

In this work, the purified pigments from different *Pseudomonas* strains underwent structural analysis, including UV, FTIR, and ^1^H/^13^C NMR spectroscopies. UV-Vis spectra revealed similar patterns in mutant model strains (PAO1 *hmgA*\* and PA14 *hmgA*\*), with dual peaks in the UVC region for PAO1 and PA14 mutants, while PAM and PexM displayed spectra indicative of melanins with more extended conjugated systems. FTIR spectra exhibited consistent features across all strains, with variations in the intensity ratios at specific bands associated to oxygenated structures. ^1^H NMR spectra showed broad signals indicative of polymeric samples, with characteristic resonances at 6.4-7.2 ppm assigned to aromatic protons. ^13^C NMR revealed carbonyl resonances, weak signals at 140-160 ppm for quaternary C-O, and intense resonances at 115-145 ppm for phenolic C-H. PAO1 *hmgA\** and PA14 *hmgA\** displayed signals at 60-110 ppm, suggesting carbohydrate moieties. Sharp resonances indicated heterocycles and unsaturated groups covalently linked to the polymer. All the spectroscopic results of PexM indicated a low content of aliphatic alcohol and ether moieties, while the high abundance of Ar units could allow a more extended conjugated system and a variety of oxidation states of the monomeric units. On the other hand, PAM strain indicated a low content of aromatic units yet with an extended conjugated structure. According to all the spectroscopic features found for the studied pyomelanin samples, a single linear polymer backbone would not support the chemical structure. Instead, we propose some of the representative monomeric units that could constitute the main polymeric backbone (Fig. 7). The combination of these units together with the substitution at phenolic or carboxylic groups would lead to a complex crosslinked polymeric structure of pyomelanin.

**Figure 7.**
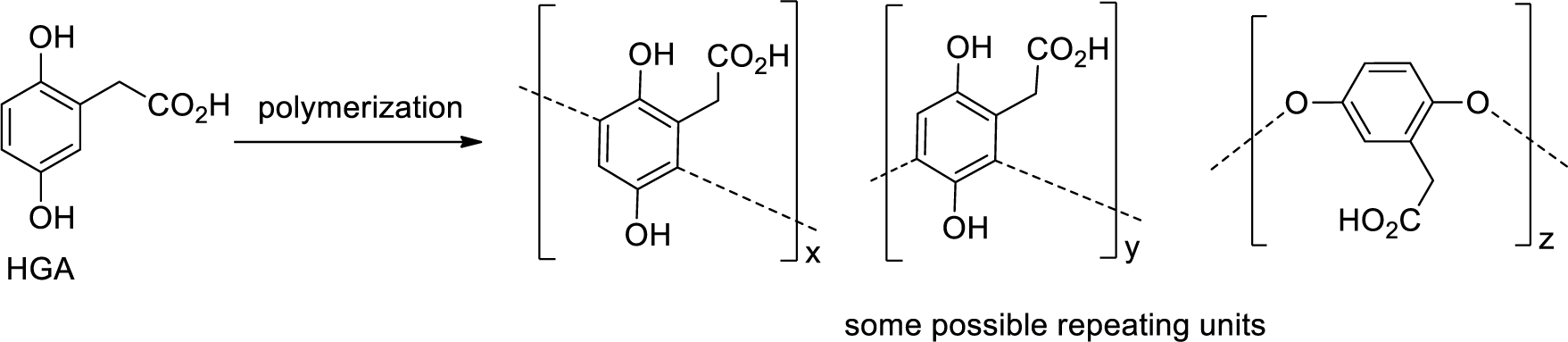
Proposed monomeric units that may constitute the complex pyomelanin structure after polymerization from homogentisate (HGA).

Melanin has been acknowledged for its advantageous attributes against UV radiation, as it functions both as an UV radiation filter and as a scavenger of reactive oxygen species (ROS). UV radiation encompasses UVA (320–380 nm), UVB (280–320 nm), and UVC (180–280 nm), with UVC possessing the highest energy but being largely absorbed by the stratospheric ozone layer (20). The UV protective characteristics of melanin have been identified in various bacterial species. Notably, melanogenic strains of *Streptomyces griseus* and *Bacillus anthracis* exhibited greater resistance to UVC and UVB radiation, respectively, than their non-melanogenic counterparts (21,22). In a melanogenic strain of *B. thuringiensis*, the observed pigment provided protection against UVC radiation for an insecticidal protein, enhancing its appeal for biotechnological applications (Liu et al., 2013). Remarkably, in certain pathogenic species, melanin exhibited effective scavenging of ROS in human fibroblasts after UVA radiation and in mouse fibroblasts after UVB exposure, concurrently increasing fibroblast viability (23,24). However, many studies are focused on analyzing the impact of UVA and UVB radiation, emphasizing the role of melanin as sunscreens (17).

In our study, we propose an examination of UVC radiation, which differs significantly from UVA and UVB due to the mechanism of its germicidal effect, mediated by chemical modifications of DNA that result from the direct action of the radiation. Our investigation revealed that pyomelanin provided protection against UVC radiation in all tested bacterial strains. Moreover, considering structural disparities observed, the addition of melanin differentially protected against UVC radiation depending on the melanin-producing strain and according with the absorption at 254nm. A pivotal experiment in this study involved assessing the survival of *P. extremaustralis* with the addition of various analyzed melanins being the PAO1’s pigment the most protective in our experimental setup.

UVC has been used since the 90’s for water disinfection, since within the non-thermal physical techniques, UVC disinfection stands out as a financially viable and readily method for potable water (25). Furthermore, in recent years, globally, technologies involving household water purifier devices employing cartridge filtration and UV disinfection have gained widespread usage (26). Given the significance of having water absent of opportunistic pathogenic bacteria such as *P. aeruginosa*, our findings regarding the protective role of melanin against UVC irradiation assume relevance.

In *P. aeruginosa*, the production of pyomelanin was first described in brown pigment clinical isolates (2). In clinical isolates from the cystic fibrosis (CF) environment, pyomelanin production has been recognized among the strategies to increase fitness in this species that enhance the virulence mainly by reducing the susceptibility to host defenses (8,27). During infection, pigmented mutants emerge with improved survival due to the capability to cope with host reactive oxygen species (28). These spontaneous pyomelanin-producing mutants account for 13% of the isolated strains in chronically infected CF patients and entail the deletion of genomic regions that lead to homogentisate accumulation (15). In this work, we investigated two pigmented mutants of the highly studied reference strains, PAO1 and PA14, affected only in *hmgA*, and PAM the natural melanogenic mutant isolated from the lungs of a patient with a pulmonary infection (not chronically infected) that suffered severe asthma. We have showed for first time melanin production of *hmgA*\* *P. aeruginosa* PAO1 and PA14 and in PAM in sputum medium. This medium mimic the nutritional and other physical characteristics, such as density of the sputum of CF patients and contains all the aminoacids (including tyrosine that can be used to produce melanin), stomach pig mucins, DNA, and lipids (29). Additionally, *P. aeruginosa* strains showed similar growth and melanin production in sputum medium in comparison with LB. In addition, the spectra of the pyomelanin obtained from PA14 *hmgA*\* strain grown in ASM gave similar polymeric signals in the aromatic and aliphatic regions of the NMR spectra than that observed in LB. Our results are relevant in the infection context since showed the bacterial capability to produce melanin in this nutritional environment besides the specific strains.

In addition, to compare with a very different species, we also analyzed a melanogenic strain (PexM) of the extremophile bacterium *P. extremaustralis* 14-3b obtained by random mutagenesis. Curiously, the transposon was inserted in a gene encoding a putative diguanylate cyclase, not previously related to melanin production. In a melanogenic isolate of *P. aeruginosa* random mutagenesis allowed the identification of genetic determinants that affected pyomelanin synthesis such as the ABC transporter, *hatABCDE* and other genes not typically associated to melanin production (30).

Moreover, an analysis of the expression of transcription factors after exposure to surface acoustic waves in *P. aeruginosa*, showed that the regulator SawR increased melanin formation through an unclear mechanism (31). There are multiple and diverse factors that affect melanin synthesis, and in different bacteria specific and general regulators such as OxyR, RpoS has been also reported as involved in melanin production (4). This suggests that diguanylate cyclase encoding by PE143B_0105645 in *P. extremaustralis* could affect homogentisate production by a direct or indirect unknown mechanism. In all analyzed strains we demonstrated that the addition of bicyclopyrone an herbicide of the class of triketones abolished pigment production, indicating the inhibition of HppD and the involvement of the homogentisate pathway in melanin production. These results are especially relevant for PAM whom genome has not been sequenced yet and the same applies to PExM for which the affected gene had not been reported as related to the homogentisate pathway.

Together, these findings underscore the considerable diversity inherent in pyomelanins from *Pseudomonas* species, highlighting discernible subgroups based on structural variations among strains, that show non-uniform grouping even within the same species. These results not only sheds light on the distinct physiological effects but also points up their relevance across various human activities.

## Material and Methods

### Bacterial strains

Different *Pseudomonas* strains were selected for this study. Model *P. aeruginosa* strains PAO1 and PA14, and its derivatives *hmgA* mutants constructed for this study (Table 3). Additionally, a spontaneous melanogenic *P. aeruginosa* strain (PAM) isolated from an Argentinean patient with chronic lung disease and a melanin-producing mutant (PexM) of *P. extremaustralis* 14-3b, an extremophile species isolated from Antarctica (32) were analyzed. Strains and plasmids employed in this study are described in Table 3.

**Table 3.** Bacterial strain and plasmids used in this study.

| Strains | Characteristics | Reference |
| --- | --- | --- |
| *P. aeruginosa* PAO1 | Wild type, wound isolate, human opportunistic pathogen | (40) |
| PAO1 *hmgA* * | Brown pigment producer, *hmgA* CRISPR-Cas isogenic mutant of PAO1 | This work |
| *P. aeruginosa* PA14 | Wild type isolate from a burn wound, highly virulent strain | (41) |
| PA14 *hmgA* * | Brown pigment producer, *hmgA* CRISPR-Cas isogenic mutant of PA14 | This work |
| *P. aeruginosa* PAM | Spontaneous brown pigment producer, isolated from a patient with obstructive pulmonary disease | This work |
| *P. extremaustralis* 14-3b DSM 25547 | Wild type, extremophile | (42) |
| *P. extremaustralis* PexM | Brown pigment producer, mini Tn5 mutant | This work |
| Plasmids |  |  |
| pMBEC6 | CRISPR/nCas9 system | (36) |
| pMBEC6(*hmgA* *PA C1205 C1206) | CRISPR/nCas9 system with the spacer to edit *hmgA* inserted | This work |
| pEX128 | Accessory plasmid of the CRISPR/nCas9 system to generate the spacer | (36) |

The PexM mutant strain was obtained during the construction of a transposon mutant library of *P. extremaustralis* (33) using pUTmini-Tn5 [ΩTc] and *E. coli* S17-1 as donor strain in a conjugation assay (34). To identify the interrupted gene, a two-step PCR strategy was performed using the following oligonucleotides: ARB1 (5’GGCCACGCGTCGACTAGTCANNNNNGATAT3’) and TN1(5’GCCCGGCAGTACCGGCATAA3’) for the first step and ARB2 (5’GGCCACGCGTCGACTAGTAC3’) and TN2 (5’GGGTGACGGTGCCGAGGATG3’) for the second step as previously described (35). The final PCR product was purified and sequenced (Macrogen, Korea).

The *P. aeruginosa hmgA* mutants of PAO1 and PA14 strains were obtained through the introduction of a stop codon by the change C1205T and C1206T in both species through a CRISPR/nCas9 system (36). For such, the spacer 5’GGGTCCCAGCGCAGGCGGTT3’ was inserted in the system and the original protocol was followed to obtain the pyomelanin-producing *P. aeruginosa* reference mutants.

### Culture conditions

Growth was analyzed using triplicates overnight precultures performed in LB medium under aerobic conditions (1:10 culture to erlenmeyer flask volume and 150 rpm agitation) at 30°C or 37°C for *P. extremaustralis* and *P. aeruginosa* strains, respectively. After centrifugation and cell washing with 0.8% saline solution to eliminate the melanin from the preculture, cells were resuspended in the same medium and used to inoculate cultures adjusted to an initial OD_600nm_ of 0.025. Cell growth was followed by plating appropriate dilutions in LB agar at different times. Plates were incubated at 37 °C for *P. aeruginosa* strains and 30 °C for *P. extremaustralis* for 24 h and colony forming units (CFU) were counted.

In addition, pigment production and cell growth were analyzed in artificial sputum medium (ASM) (29) after 24h growth for *P. aeruginosa* strains.

### Pigment purification

After 48 h of incubation, 100 mL of cell-free culture supernatants were acidified (pH 2) using HCl 5 M and incubated at room temperature in the dark for 48 h. Then, the precipitated pyomelanin was centrifuged and washed twice with milli Q quality water, once with an acetone:ethanol solution (1:1) and resuspended in water at 100°C for 15 min. After centrifugation, the precipitate was washed twice with water and freeze dried to give purified pyomelanin that was stored at -20°C.

### Structural analysis

An FT-IR (ATR) analysis was performed in a Thermo Scientific Nicolet IS50 FTIR spectrometer. For the UV-Vis absorbance, 1 mg. mL^-1^ of purified pyomelanin was solubilized in 0.2 M NaOH and was further diluted to 0.05 mg. mL^-1^ and the spectra was measured in a Shimadzu UV-2600i spectrophotometer, using quartz cells of 1 cm optical path length. Finally, NMR spectra of pyomelanin (15 mg), dissolved in NaOD-D_2_O (0.6 mL), were recorded with a Bruker Avance Neo 500 instrument (^1^H: 500 MHz, ^13^C 125.4 MHz) with residual solvent as the internal standard.

### Pigment synthesis inhibition

The specific inhibitor bicyclopyrone[4-hydroxy-3-(2-(2-methoxy-ethoxymethyl)-6-trifluoromethyl-pyridine-3-carbonyl)-bicyclo (3,2,1) oct-3-en-2-one] (ACURON^TM^ UNO, Syngenta Agro S.A.) was used to analyze the relevance of HppD. Different concentrations (0.1 mM, 1 mM, and 10 mM) of this inhibitor were added to test tubes cultures performed in LB medium as previously described. Bacterial growth and culture coloration were recorded to analyze the pyomelanin production.

### UVC survival assay and decay modeling

UVC (254 nm) tolerance in wild-type and pyomelanin-producing mutant cells was evaluated by irradiating cells obtained from LB overnight cultures. The cells, washed and adjusted to 0.3 OD_600nm_ in PBS, were divided into two 10 mL samples: one with 0.15 mg mL^-1^ purified pyomelanin and the other without. Both samples were exposed to a fluence rate of 0.93 W.m^-2^ using a low-pressure mercury lamp (Phillips TUV 15W/G15T8) encased in a pseudo-collimated bean device (37) within 135 and 360 s for samples without or with melanin, respectively. Fluence rate was measured with an ILT77 Germicidal Radiometer equipped with a XP77CE sensor (International Light Technology), following the procedures described by Bolton and Linden (2003)(38). Viable cell counts were measured by plating in LB, followed by dark incubation. Survival, expressed as a logarithmic fraction of CFU.mL^-1^ at initial time (N_0_), showed increasing slopes initially, decreasing suddenly beyond a certain fluence, and remaining constant at high fluences. Biphasic curves suggest inactivation of a majority susceptible subpopulation initially and presence of a minority highly tolerant subpopulation later. Biphasic survival curves were described assuming the presence of two bacterial subpopulations, by adjusting to the experimental results the parameters in the equation:

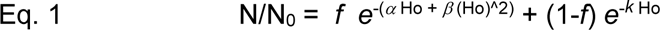

In the equation, N/ N_0_ signifies survival after fraction H_0_, with f as the fraction for the majority subpopulation. The first term, using the linear-quadratic model (39), addresses majority subpopulation inactivation. The second term depicts tolerant subpopulation inactivation as exponential decay, with k as the inverse of fluence reducing survival by 1/e (0.37).

Attenuation of the incoming radiation due to absorption by pyomelanin was estimated from the absorbance of the irradiation medium (PBS) and the depth of the samples, using an equation derived from the integration of the expression of the Lambert-Beer law (38).Results were expressed as percentage.

### Statistical analysis

The significance of the differences between means in the different conditions were evaluated through non-parametric one-way analysis of variance (ANOVA) with a Kruskal-Wallis test with multiple comparisons, and unpaired and non-parametric t-test Mann-Whitney correction, with confidence levels at > 95% (i.e., P < 0.05 was considered as significant). All the analyses were performed using GraphPad Prism version 8.0.2 for Windows.

## Acknowledgments

This work was funded by grants from Universidad de Buenos Aires 20020220400198BA and 20020220300089BA, CONICET 1120210100680CO and Agencia Nacional de Promoción de la Investigación, el Desarrollo Tecnológico y la Innovación (Agencia I+D+i) PICT 2019-02482 and PICT-2021-GRFTI-00265. A.K., N.I.L and P.M.T. are career investigators from CONICET. M.N.D.A. hold a doctoral fellowship from Agencia I+D+i.

## Ethical statements

The PAM strain was collected as part of the project approved by the Ethical Committee of the “Hospital General de Niños Dr. Pedro Elizalde” under the approval number PRIISA BA: 4466. All patients signed the information and consent forms.

