## Supplementary Figures for "BEYOND UNIFORMITY: Pyomelanin’s structural complexity impacts on UV shielding in *Pseudomonas* species with different lifestyles"

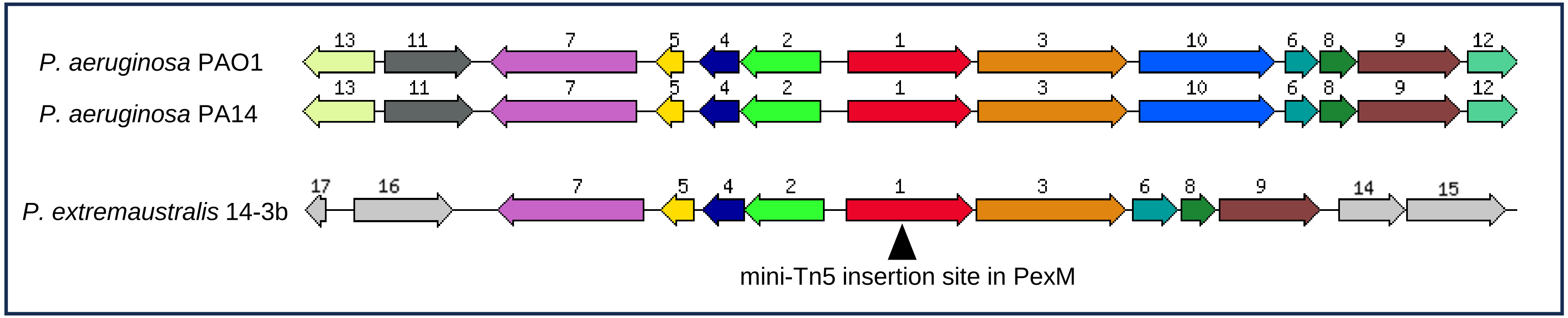


| Reference | Encoding product |
|---|---|
| 1 | Probable two-component response regulator |
| 2 | Uncharacterized protein conserved in bacteria |
| 3 | Methyl-accepting chemotaxis protein |
| 4 | NTP pyrophosphohydrolases including oxidative damage repair enzymes |
| 5 | Hypothetical protein |
| 6 | 3-dehydroquinate dehydratase II |
| 7 | Biosynthetic arginine decarboxylase |
| 8 | Biotin carboxyl carrier protein of acetyl-CoA carboxylase |
| 9 | Biotin carboxylase of acetyl-CoA carboxylase |
| 10 | Cytochrome c-type biogenesis protein DsbD, protein-disulfide reductase |
| 11 | Hypothetical protein |
| 12 | Hypothetical protein |
| 13 | Ferrichrome-iron receptor |
| 14 | Ribosomal protein L11 methyltransferase |
| 15 | Probable transmembrane protein |
| 16 | DNA-damage-inducible protein F |
| 17 | Secreted protein Hcp |

Fig. S1. Comparative analysis of the genomic region of *P. aeruginosa* PAO1 and PA14, and *P. extremaustralis* 14-3b showing mini-Tn5 insertion site in PexM. Locus_tag in PexM (PE143B_0105645). Region size: 16,000 bp. The same color indicates homologous genes.

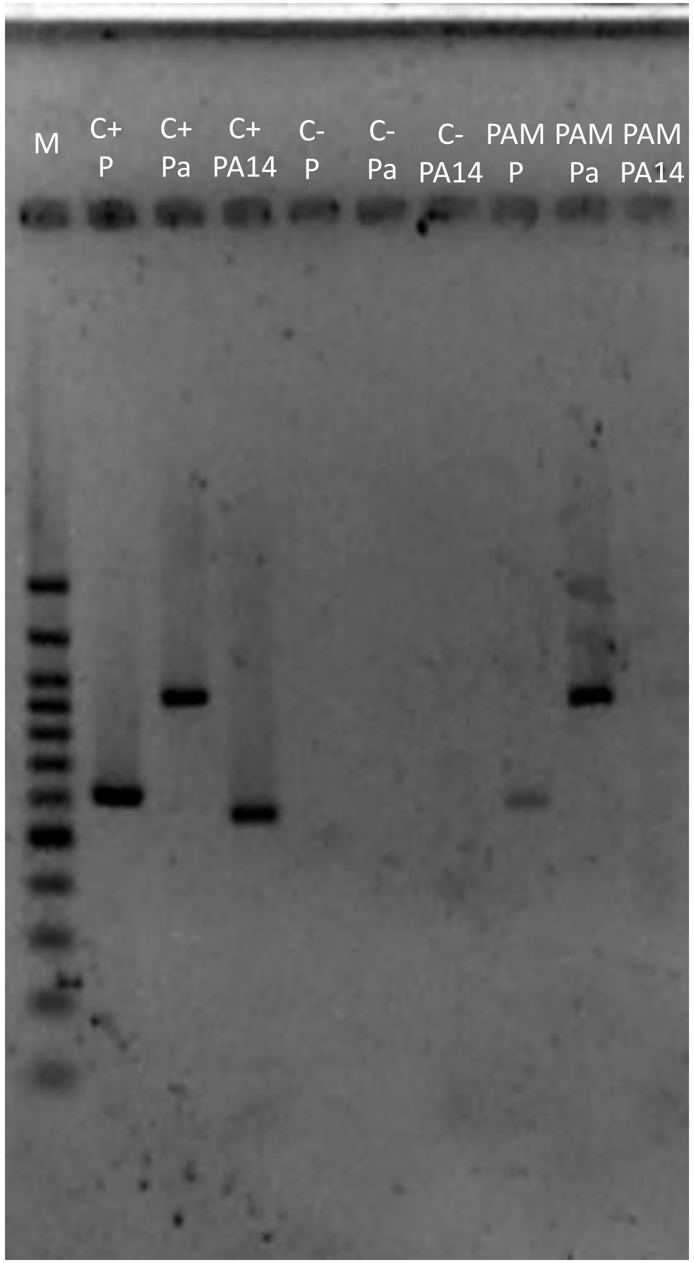


Fig. S2: Identification of PAM strain through specific PCR. For *Pseudomonas* genus, “P” primer set ( 5'GACGGGTGAGTAATGCCTA3' and 5'CACTGGTGTTCCTTCCTATA3’) was used and for *P. aeruginosa*, “Pa” primers (5'GGGGGATCTTCGGACCTCA3' and 5'TCCTTAGAGTGCCCACCCG3’). These primers were taken from Spilker et al. (2003).
For PA14 strain ("PA14") specific primers 5'TCGACCTTCAGTGTTTCCCG3' and 5'ACGGTG TGATGGGCAATGAA3’ were designed using the Primer BLAST tool from the NCBI to amplify a fragment of *ybtQ* (GenBank: AY049068.1) gene specific of PA14 (Choi et al. 2002). Lines: M, Marker 100 bp (); Positive controls (C+): P, *P. extremaustralis;* Pa, *P. aeruginosa* PAO1, and PA14: *P. aeruginosa* PA14; Negative controls ("C-"): P, *Clostridium sp.;* Pa, *P. extremaustralis*, and PA14, *P. aeruginosa* PAO1; PAM P, PAM Pa, and PAM PA14.

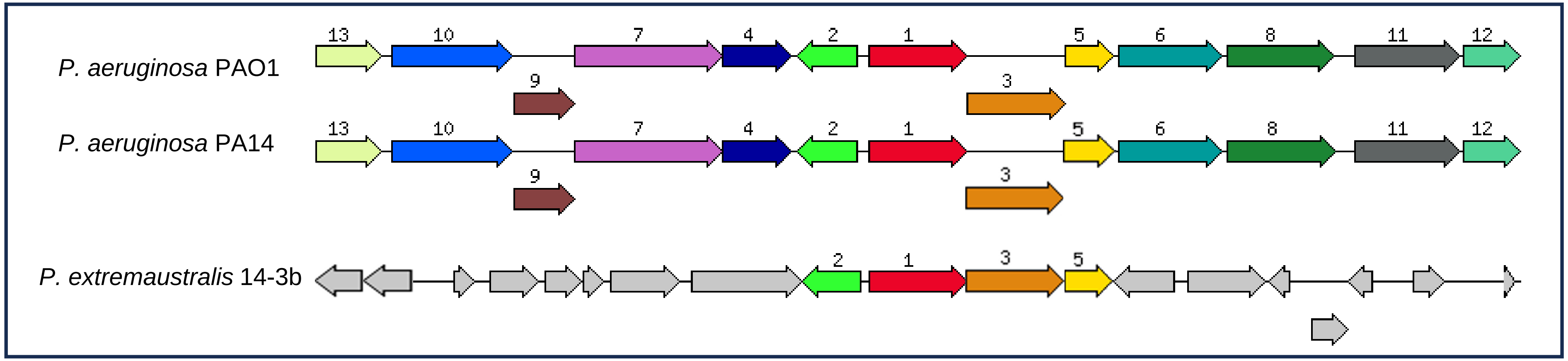


| Reference | Encoding product |
|---|---|
| 1 | Homogentisate 1,2-dioxygenase |
| 2 | Transcriptional regulator, IclR family |
| 3 | Fumarylacetoacetase |
| 4 | Hydroxymethylglutaryl-CoA lyase |
| 5 | Maleylacetoacetate isomerase |
| 6 | Probable MFS transporter |
| 7 | Methylcrotonyl-CoA carboxylase biotin-containing subuni |
| 8 | probable transcriptional regulator |
| 9 | Methylglutaconyl-CoA hydratase |
| 10 | Methylcrotonyl-CoA carboxylase carboxyl transferase subunit |
| 11 | D-beta-hydroxybutyrate permease |
| 12 | D-beta-hydroxybutyrate dehydrogenase |
| 13 | Isovaleryl-CoA dehydrogenase;Butyryl-CoA dehydrogenase |

Fig. S3. Comparison of genomic region containing *hmgA* in *P. aeruginosa* PAO1 and PA14, and *P. extremaustralis* 14-3b. Region size :16,000 bp. The same color indicates homologous genes.

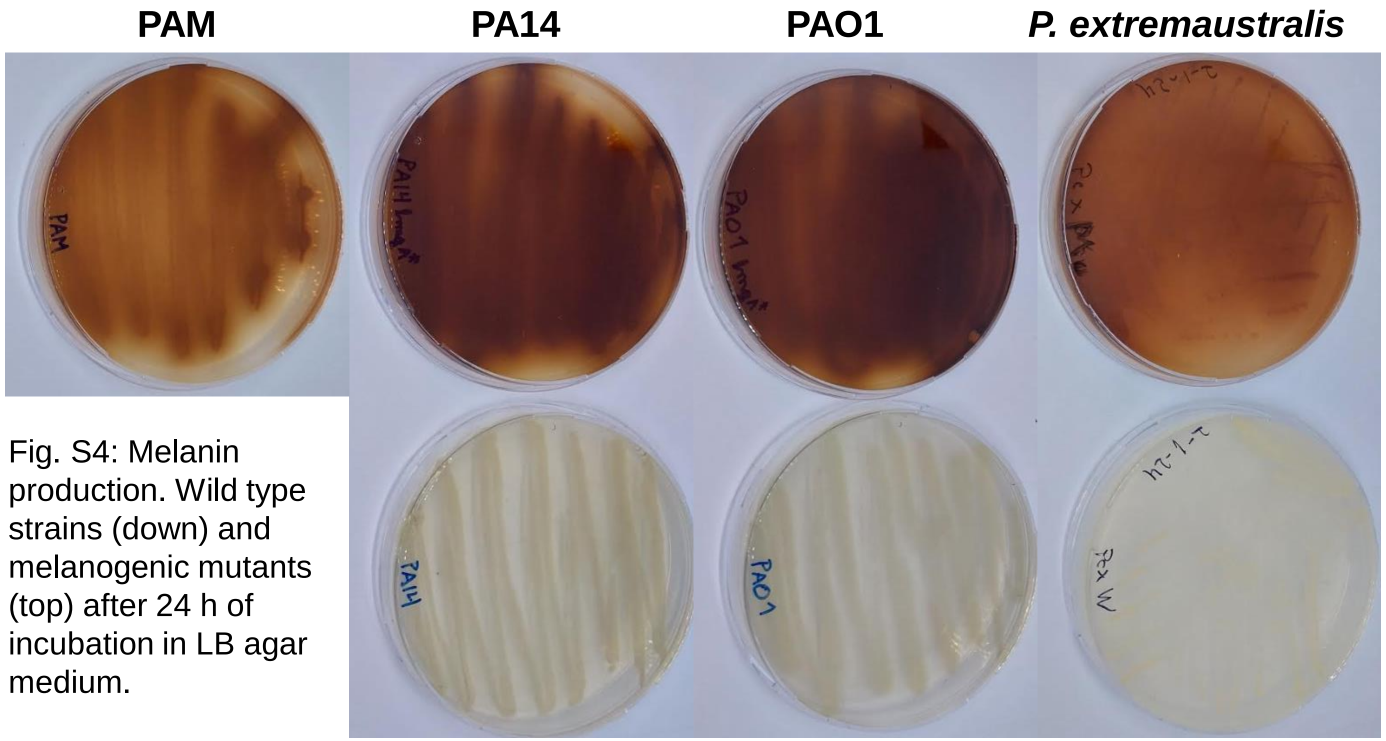


Fig. S4: Melanin production. Wild type strains (down) and melanogenic mutants (top) after 24 h of incubation in LB agar medium.

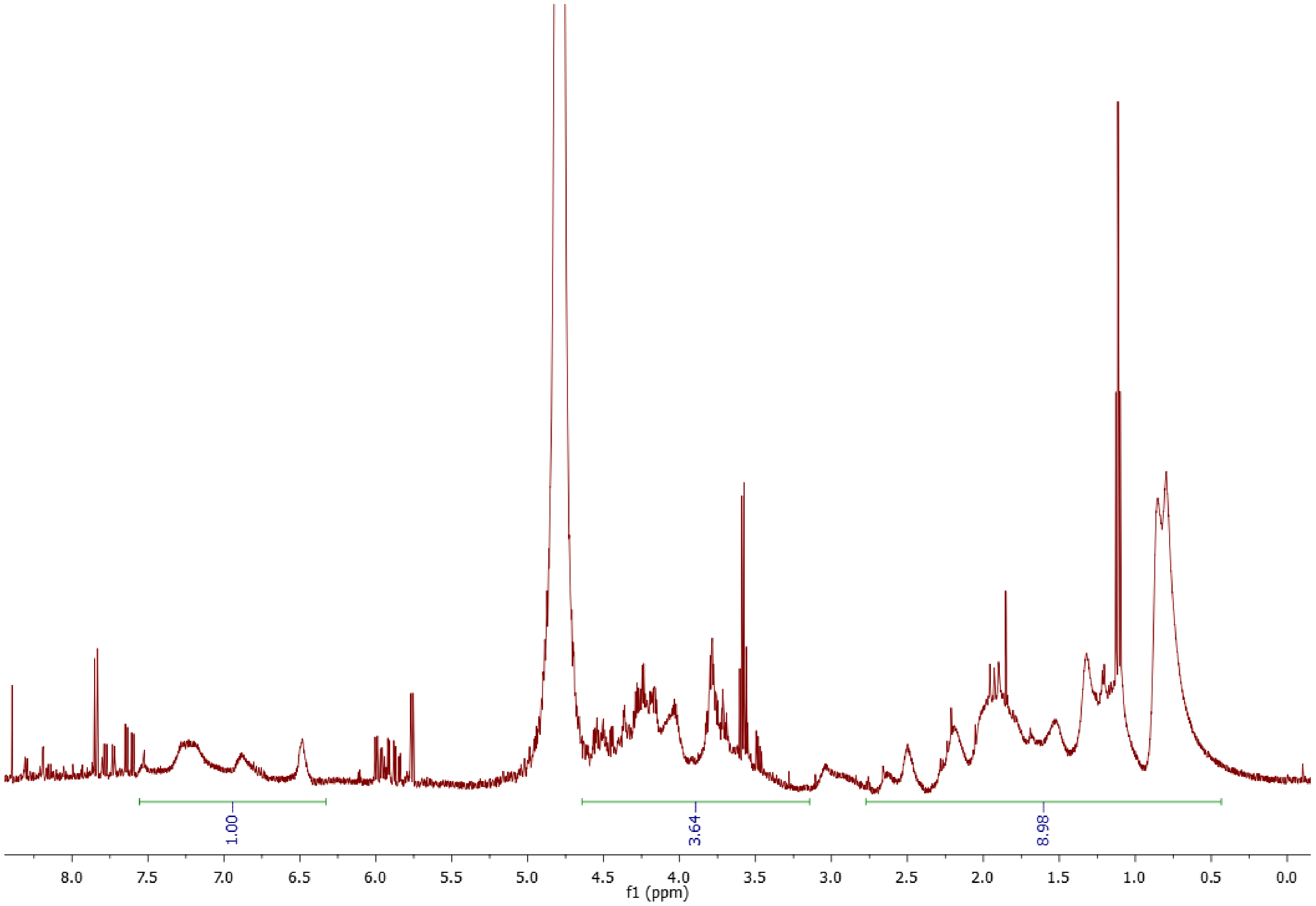


Fig. S5: Integrated $^{1}H$ NMR spectra of purified PAO1 *hmgA** pyomelanin. Cells were grown in LB medium.

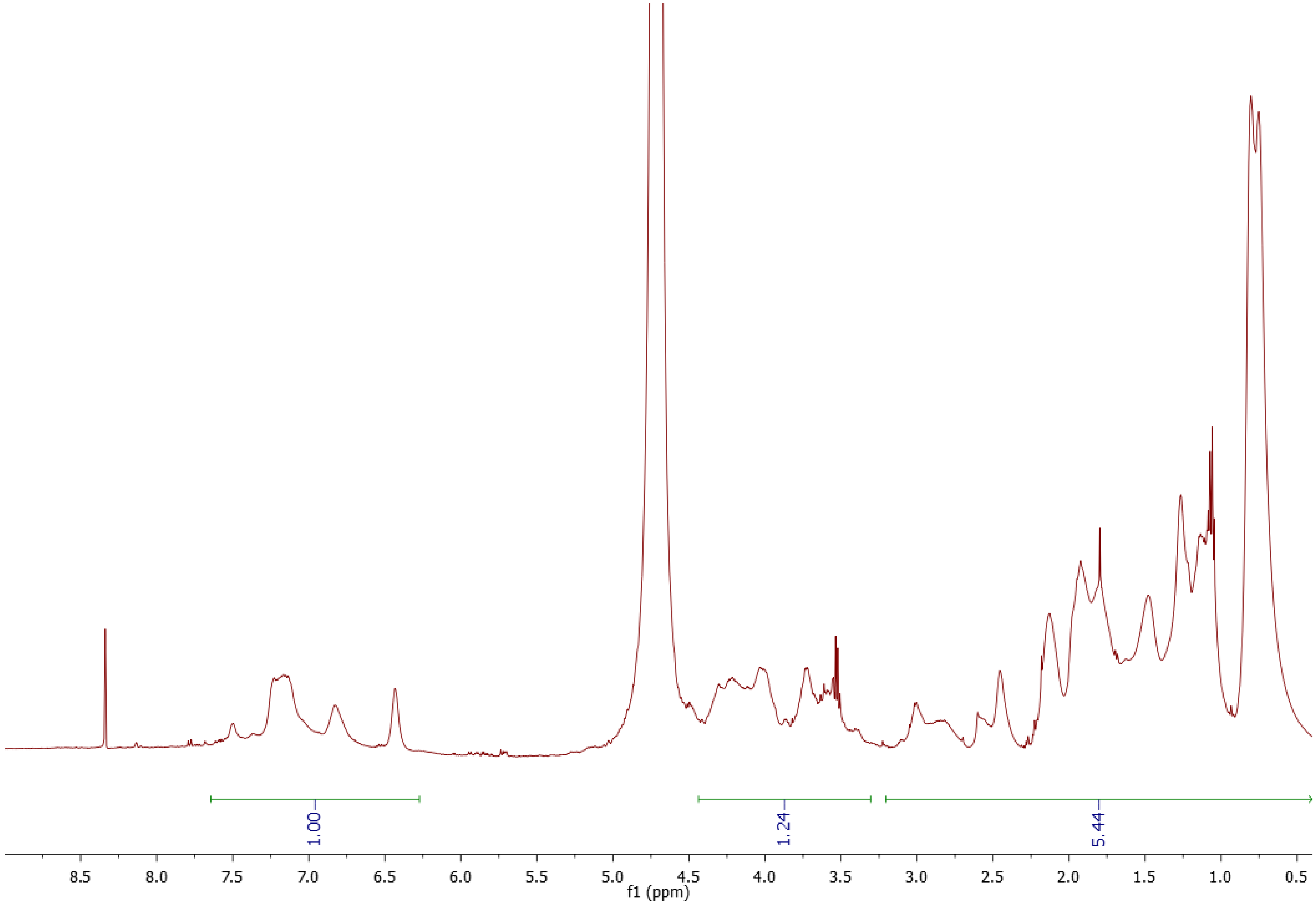


Fig. S6: Integrated $^{1}H$ NMR spectra of purified PexM pyomelanin. Cells were grown in LB medium.

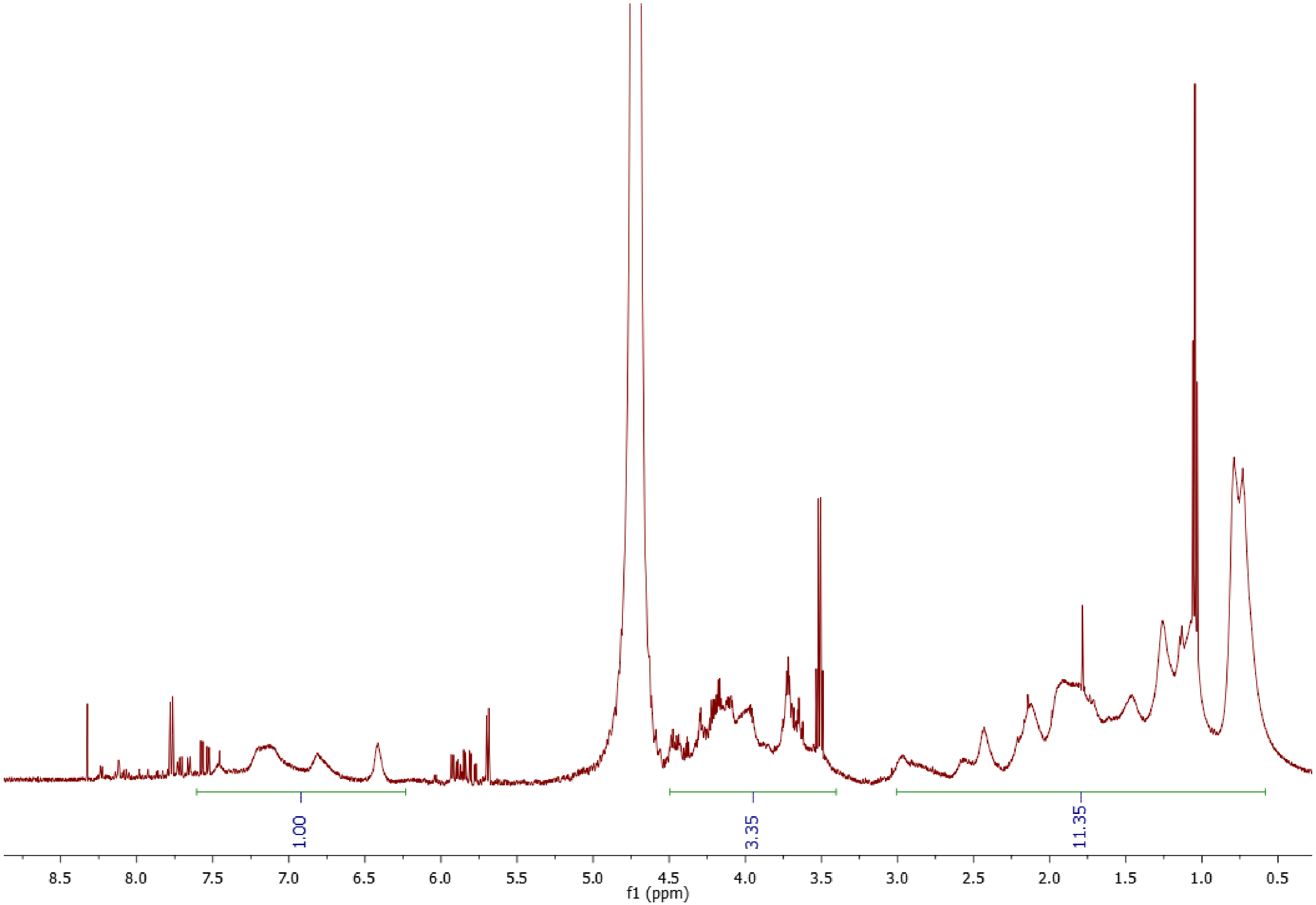


Fig. S7: Integrated $^{1}H$ NMR spectra of purified PAM pyomelanin. Cells were grown in LB medium.

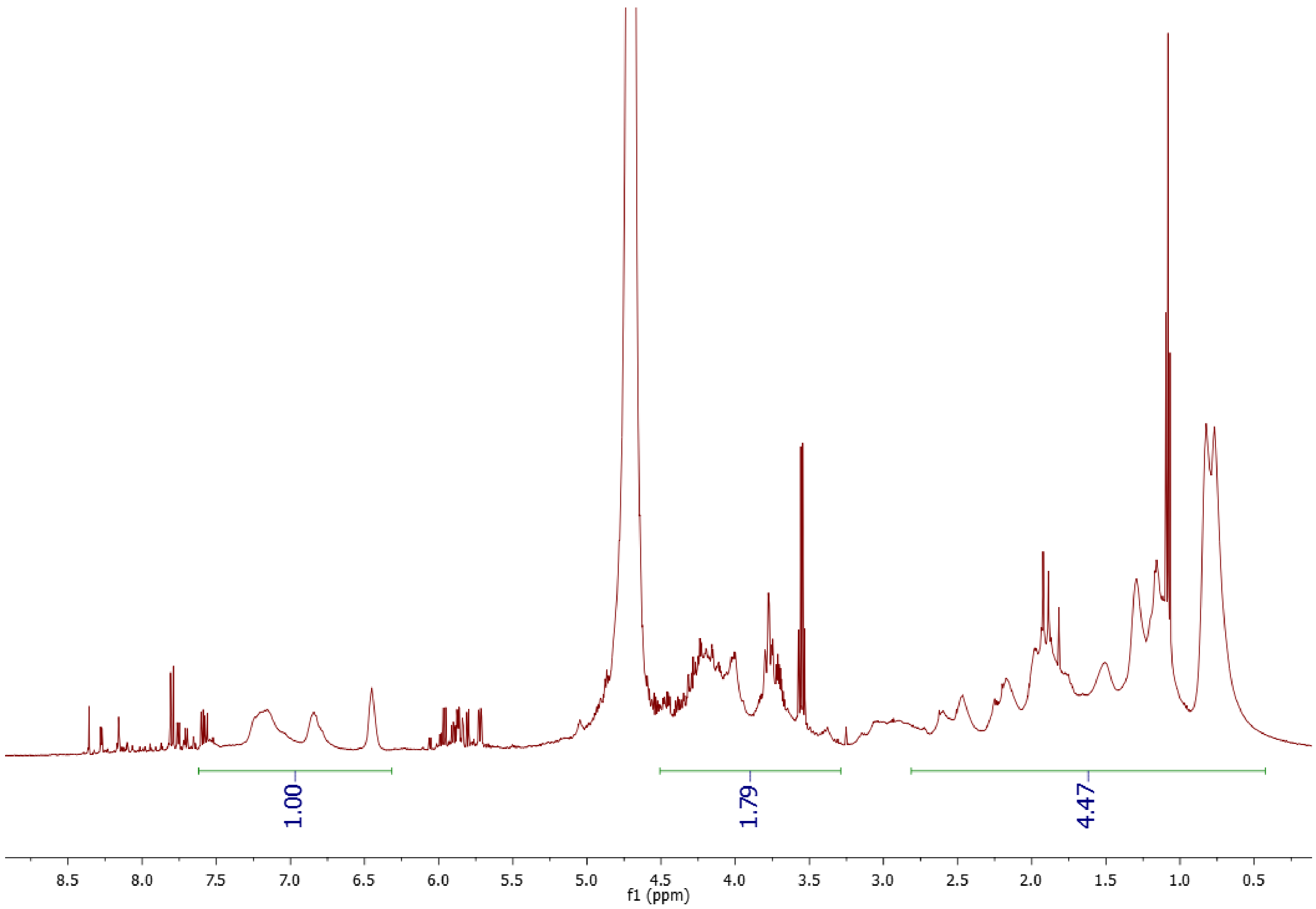


Fig. S8: Integrated $^{1}$H NMR spectra of purified PA14 *hmgA** pyomelanin. Cells were grown in LB medium.

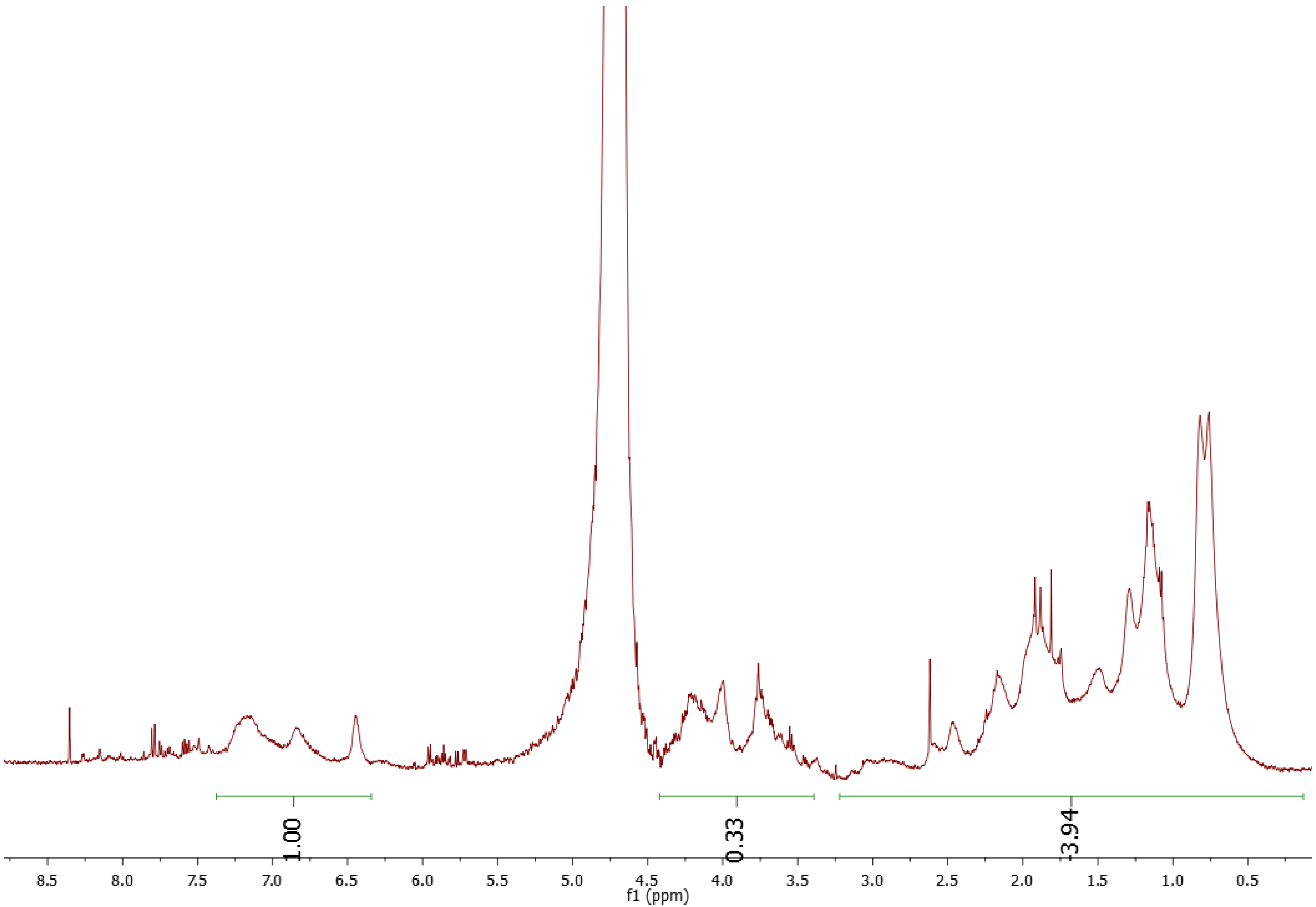


Fig. S9: Integrated $^{1}H$ NMR spectra of purified PA14 *hmgA** pyomelanin. Cells were grown in ASM medium.

| Strain | Survival modeling |
|---|---|
| PAO1 | log(0.993314406*(2.718^(-1*(0.02731*D(T)+0.00167*D(T)^2)))+(1-0.993314406)*(2.718^(-0.0398*colD(T)))) |
| PAO1 suplemented with PAO1 *hmgA**'s melanin | log(0.995322002*(2.718^(-1*(0.00248*D(T)+0.00037*D(T)^2)))+(1-0.995322002)*(2.718^(-0.0126*colD(T)))) |
| PAO1 *hmgA** | log(0.9999652*(2.718^(-1*(0.05412*D(T)+0.00052*D(T)^2)))+(1-0.9999652)*(2.718^(-0.01077609824*D(T)))) |
| PAO1 *hmgA* suplemented with PAO1 *hmgA**'s melanin | log(0.997230891*(2.718^(-1*(0.01223*D(T)+0.00012*D(T)^2)))+(1-0.997230891)*(2.718^(-0.0092*D(T)))) |
| PA14 | log(0.9998381845*(2.718^(1*(0.0633*D(T)+0.00027*D(T)^2)))+(1-0.9998381845)*(2.718^(-0.0032006*D(T)))) |
| PA14 suplemented with PA14 *hmgA**'s melanin | log(0.9864190547*(2.718^(-1*(0.02094*D(T)+0.0001*D(T)^2)))+(1-0.9864190547)*(2.718^(-0.01819042223*D(T)))) |
| PA14 *hmgA** | log(0.9950781514*(2.718^(-1*(0.05389*D(T)+0.00019*D(T)^2)))+(1-0.9950781514)*(2.718^(-0.0221969203*D(T)))) |
| PA14 *hmgA* suplemented with PA14 *hmgA**'s melanin | log(0.9887539503*(2.718^(-1*(0.01718*D(T)+0.00005*D(T)^2)))+(1-0.9887539503)*(2.718^(-0.01441418268*D(T)))) |
| *P. extremaustralis* | log(0.9998691029*(2.718^(-1*(0.02448*D(T)+0.00097*D(T)^2)))+(1-0.9998691029)*(2.718^(-0.008657719*D(T)))) |
| *P. extremaustralis* suplemented with PexM's melanin | log(0.9996395957*(2.718^(-1*(0.00.00237*D(T)+0.00023*D(T)^2)))+(1-0.9996395957)*(2.718^(-0.0075064*D(T)))) |
| PexM | log(0.999684*(2.718^(-1*(0.05389*D(T)+0.00019*D(T)^2)))+(1-0.999684)*(2.718^(-0.0005*D(T)))) |
| PexM suplemented with PexM's melanin | log(0.999957702*(2.718^(-1*(0.00849*1.868*D(T)+0.00021*3.49*D(T)^2)))+(1-0.999957702)*(2.718^(-0.00396*D(T)))) |
| PAM | log(0.9998443138*(2.718^(1*(0.05588*0.45387*D(T)+0.00055*0.206*D(T)^2)))+(1-0.9998443138)*(2.718^(-0.0025789*0.45387*D(T)))) |
| PAM suplemented with PAM's melanin | log(0.9994480115*(2.718^(-1*(0.01783*D(T)+0.00014*D(T)^2)))+(1-0.9994480115)*(2.718^(-0.00872697*D(T)))) |

Table S1: Function modeling corresponding to each strain survival in presence or absence of melanin in the solution exposed to UVC. D(T) is the time of exposure.
